# Targeting host eIF4A with zotatifin inhibits Bundibugyo virus and other filoviruses

**DOI:** 10.64898/2026.11.18.767653

**Authors:** Katarzyna Bogucka-Janczi, Hajime Yurugi, Nadine Biedenkopf, Franziska Kraft, Olena Shtanko, Vladimir Gonçalves Magalhães, Sandra Ciesek, Zhuoyi Song, Davide Ruggero, Victor Moyo, Tawanda Gumbo, Krishnaraj Rajalingam

## Abstract

Filoviruses, including Ebola and Marburg viruses, depend on host eIF4A helicase activity for replication. Zotatifin is a first-in-class, clinical-stage eIF4A inhibitor that acts as a molecular glue, stabilizing the interaction between eIF4A and selected polypurine-rich RNA sequences, thereby inhibiting RNA unwinding and ribosomal scanning. Here, we identify pronounced polypurine enrichment in the 5′ untranslated regions (5′ UTRs) of the seven major filoviral protein-coding genes NP, VP35, VP40, GP, VP30, VP24, and L across four virus species, including Bundibugyo virus (BDBV). Consistent with these findings, zotatifin inhibits the replication of Ebola, Marburg, Sudan, Taï Forest, and Bundibugyo viruses at low-nanomolar concentrations. Zotatifin has been administered to humans in oncology and COVID-19 clinical studies. Clinical pharmacokinetic–pharmacodynamic (PK–PD) modeling indicates that exposures resulting in inhibition of BDBV replication are achievable in humans at tolerated doses following subcutaneous administration. Together, these findings support the repurposing of zotatifin as a host-directed antiviral candidate against BDBV and other filoviruses, with potential applications in preparedness for future outbreaks.

**Highlights:**

- Zotatifin targets host eIF4A to inhibit filovirus replication.
- Conserved polypurine-rich motifs in filoviral 5′ UTRs are putative sites for zotatifin-induced eIF4A clamping.
- Zotatifin suppresses EBOV, TAFV, SUDV, MARV and BDBV replication at 6.25 nM in vitro, with IC50 and IC90 values of 1.0–3.5 nM for EBOV, MARV and BDBV.
- Monte Carlo simulation with a population PK model built from COVID-19 patient data identified subcutaneous zotatifin regimens that keep concentrations above the BDBV IC□□ for ≥50% of the dosing interval in 540,000 virtual pediatric and adult subjects.
- Predicted exposures for these regimens fall within those achieved with subcutaneous doses tolerated in patients with COVID-19.

## Main text

Filoviruses, including Orthoebolavirus and Orthomarburgvirus (MARV) are non-segmented, negative-strand RNA viruses ((−)ssRNA), which cause severe hemorrhagic fevers ^1,2^. Four species within the *Orthoebolavirus* genus cause Ebola virus disease in humans: *Orthoebolavirus zairense (Ebola virus, EBOV), Orthoebolavirus sudanense (Sudan virus, SUDV), Orthoebolavirus bundibugyoense (Bundibugyo virus, BDBV), and Orthoebolavirus taiense (Taï Forest virus, TAFV)*. While Orthomarburgvirus belongs to a separate genus, it causes a clinically indistinguishable severe hemorrhagic fever (Marburg virus disease)^1^.

The ongoing BDBV outbreak in the Democratic Republic of the Congo has confirmed 10,467 cases and 5,103 deaths, a crude case fatality ratio (CFR) of 48%, as of 20 October 2026^3^. The CFR is worse among children^4^. These viruses primarily target cells of the monocyte-macrophage lineage, where viral replication drives systemic viremia. Following the initial infection, infected cells and free viral particles disseminate to lymphoid tissue, vascular organs, the liver, spleen, and adrenal glands. Hemorrhage ensues from damage to blood vessels and disseminated intravascular coagulation (DIC)^5–7^. The incubation period is 2–21□days, and in severe forms death occurs between 6 and 16□days after symptoms appear.

High genetic diversity and mutation rates among filoviruses, including EBOV and BDBV, which share only 66% sequence homology, pose a major challenge for high-precision but narrow-spectrum countermeasures. Because of this high sequence divergence, direct-acting antivirals and monoclonal antibodies frequently demonstrate virus- or species-specific activity^8^.

Filoviruses are reliant on the host cell’s canonical 7-methylguanosine (m^7^G) cap-dependent translation. Replication begins when the viral RNA-dependent RNA polymerase (RdRp) complex, led by the L protein, transcribes the (−)ssRNA into 5′-capped, monocistronic, positivesense messenger RNAs (mRNAs), including Nucleoprotein (NP), Viral proteins (VP) such as VP35 (cofactor of replication/transcription complex), VP40 (matrix protein), VP30 (cofactor of replication/transcription complex), VP24 (matrix protein), Glycoprotein (GP), and Large Protein (L-protein) (an RNA-dependent RNA polymerase) ^2,9–12^. The trimeric host translation initiation complex, eukaryotic initiation factor 4F (eIF4F), composed of the cap-binding protein eIF4E, the scaffold protein eIF4G and the DEAD-box helicase eIF4A, binds directly to the 5′ m^7^G cap structure of the mRNA template. The eIF4F complex then uses the helicase activity of eIF4A to unwind any secondary RNA loop structures in the 5′ UTR, clearing the path for host ribosome scanning and completion of protein translation ^13–18^.

Natural rocaglates such as silvestrol are potent antiviral agents targeting the host translation^14,16,17,19^. Structural studies have demonstrated that silvestrol potently suppresses luciferase reporter gene activity driven by the 5′ UTRs of several ebolavirus mRNAs^20^. Zotatifin is a synthetic rocaglate derivative optimized to overcome the structural hurdles of natural rocaglates^21^. It targets eIF4A helicase, forcing it to clamp onto highly structured, polypurine-rich (AG-rich) motifs within the 5′ UTRs of mRNAs, ultimately halting ribosome scanning and blocking viral protein synthesis^13–17,22,23^. Moreover, eIF4A inhibition by zotatifin upregulates interferon-stimulated genes (ISGs) in patient biopsies collected during a Phase 1/2 clinical trial (NCT04092673), suggesting a dual antiviral mechanism—one direct and the other host-directed^24^. Pathway-level reasoning suggests that eIF4F inhibitors are unlikely to worsen the thrombin-driven DIC characterizing ebolavirus disease.

Ebolaviruses have a conserved 12-nucleotide sequence, 5′-GAGGAAGAUUAA-3′, which serves as the precise consensus transcription start site (TSS)^11,25^. We assessed the degree of sequence conservation within the 5′ UTR polypurine-rich motifs across diverse ebolavirus mRNAs. We analyzed 5′ UTR sequences of seven major ebolavirus protein-coding genes (NP, VP35, VP40, GP, VP30, VP24, and L) across four viral species and quantified local purine (A/G) content along a 10-nucleotide sliding window. Strikingly, nearly all genes exhibited marked purine enrichment in the 5′-proximal regions, with local A/G content exceeding 70% in most sequences **(Figure 1A and S1)**.

**Figure 1.**
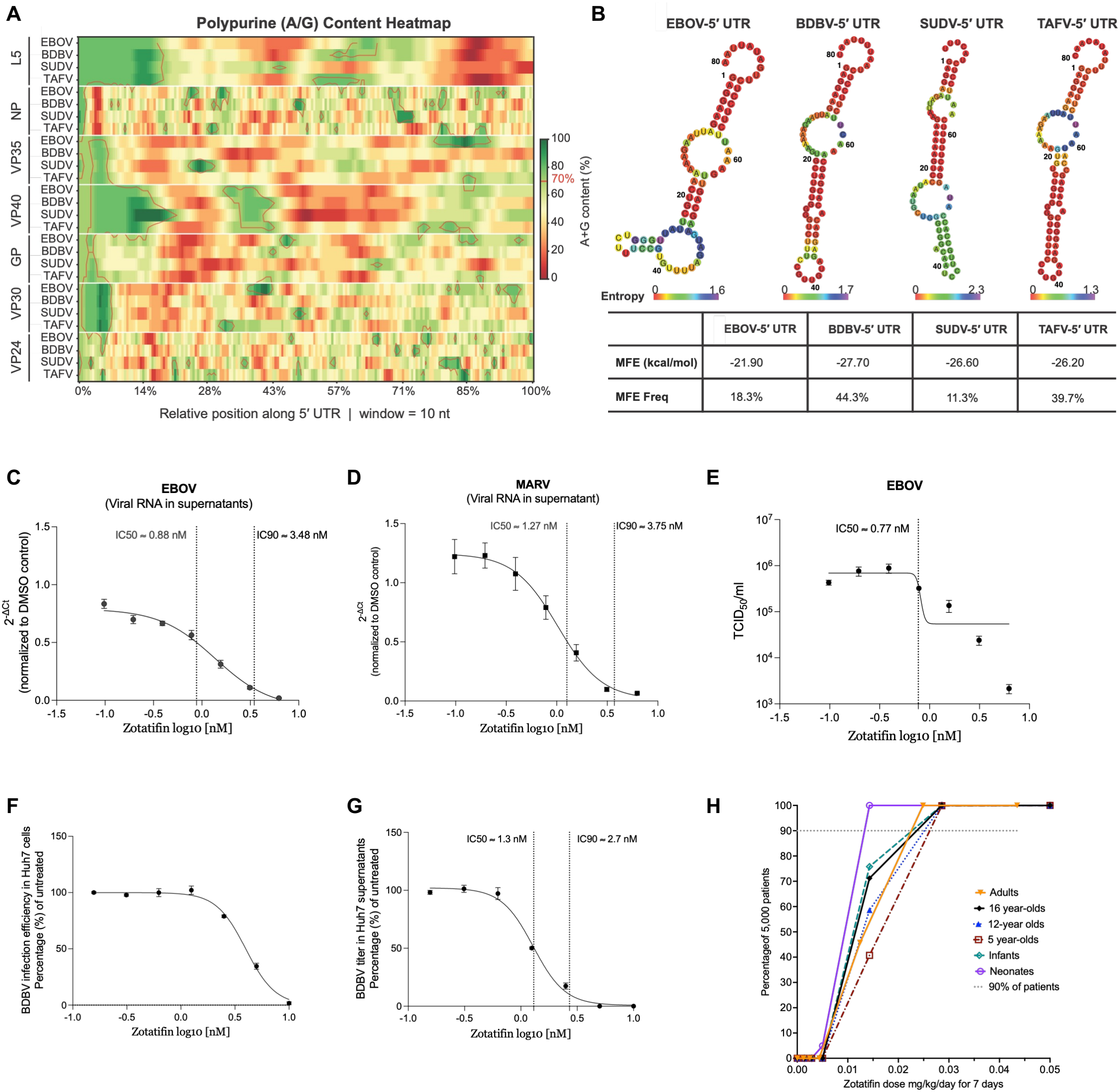
Inhibition of *Filoviridae* replication through targeted host eIF4A modulation by zotatifin. **(A–B) Purine enrichment in the 5′-proximal regions of Ebola virus mRNA 5′ UTRs is conserved across Ebola virus genes in different species. (A)** Purine (A/G) content heatmap across the 5′ UTRs of seven Ebola virus protein-coding genes in four Orthoebolavirus species. Color indicates local A+G percentage (%) within a 10-nucleotide window. Red circles mark putative polypurine tracts where local A/G content exceeds 70%. **(B)** Predicted secondary structure of the EBOV L gene 5′ UTR generated by *RNAfold*. The structure shown represents the minimum free energy (MFE) prediction, and the MFE value and frequency are presented in the table below. The color of each nucleotide indicates positional entropy: red, low entropy (well defined); green/blue, high entropy (disordered). Positions with low entropy are more likely to follow the MFE-structure prediction *in vivo*. **(C–E)** The antiviral efficacy of zotatifin against filoviruses was assessed in Huh7 cell cultures infected with the indicated filovirus strain. Briefly, cells were infected at an MOI of 0.1 as indicated. After infection, zotatifin or vehicle control (DMSO) was added to the medium at indicated final concentrations (6.25–0.1 nM, two-fold dilutions). Supernatants from infected cells were collected 48 h post-infection, RNA was isolated and subjected to qRT-PCR detecting filovirus-specific RNA in the supernatant of infected cells. Shown are 2-(dCt value, normalized to DMSO control (1). Results are presented as mean ± SEM of at least three independent experiments (**(C)** EBOV, n = 4; **(D)** MARV, n = 3). **(E) EBOV egress from zotatifin-treated Huh7 cells**. VeroE6 cells were inoculated with serial dilutions of supernatant harvested from Huh7 cells. TCID_50_/ml was calculated using the Spearman–Kärber method. Results are presented as mean ± SEM of 3 independent experiments (n = 3). **(F–G) Inhibition of BDBV replication by zotatifin**. Focus-forming assay (FFA) for Bundibugyo virus (BDBV) was used as an immunodetection-based quantitative measure of infection efficiency in cell monolayers **(F)** Cells were infected with Bundibugyo virus (BDBV; MOI = 0.1) for 2 h and treated with two-fold serial dilutions of zotatifin. After 48 h, supernatants were titrated on Vero cells **(G)**. Results are presented as mean ± SEM of 3 independent experiments. **(H)** Monte Carlo experiments cumulative response fraction. Doses (mg/kg) of 0, 0.0014, 0.003, 0.004, 0.014, 0.029, and 0.043, were administered subcutaneously once a day for 7 days. Each symbol is a mean of 5,000 virtual subjects. The acceptable dose is the lowest one above 90% proportion of patients, which was 0.025 mg/kg/day for all age groups except neonates (0.014 mg/kg/day).

To examine the structural context of these sequences more closely, we predicted the secondary structures of the 5′ UTR of each mRNA using *RNAfold*^*26*^. Among the seven monocistronic mRNAs, the L 5′ UTR formed the most thermodynamically stable structure, characterized by a high minimum free energy (MFE) frequency, low ensemble diversity, and strong concordance across different prediction models (**Figure 1B**). The predicted secondary structures of the L 5′ UTR were highly similar across all four ebolavirus species **(Figure 1B)**. Notably, although other regions of the 5′ UTR show elevated positional entropy—indicating that these regions are conformationally flexible and can adopt multiple alternative structures—the 5′-proximal nucleotides consistently exhibit a well-defined and evolutionarily conserved secondary structure with entropy near zero (**Figure 1B**). Interestingly, this pattern of 5′-proximal structural conservation was similarly observed in other genes, including VP35 **(Figure S2A)** and VP40 **(Figure S2B)**, suggesting that it represents a shared feature of Ebola virus mRNA 5′ UTRs rather than a gene-specific property.

Targeting these conserved, AG-rich eIF4A-binding motifs in the 5′ UTR of crucial ebolavirus mRNAs offers a highly effective antiviral strategy that prevents single-nucleotide mutational escape. These motifs are embedded directly within the transcription start sites (TSS) of the viral mRNAs, making them ideal therapeutic targets across ebolavirus species.

To evaluate the antiviral efficacy of zotatifin against filoviruses, we initially profiled its activity across multiple filoviral species. Zotatifin treatment resulted in a profound, dose-dependent reduction in the replication of all filoviruses tested, with almost no residual viral RNA remaining after treatment with 6.25 nM (Figure S3A; see also Figure 1C–1G).

To define zotatifin’s potency more precisely, we performed follow-up dose-response titrations using two-fold serial dilutions below the 6.25 nM threshold. To assess virus egress from zotatifin-treated Huh7 cells, cell-free supernatants from the treated Huh7 cells were collected and inoculated onto Vero cells to evaluate the production and subsequent infectivity of progeny virus (**Figure 1E, 1G)**.

These independent assays confirmed that zotatifin efficiently reduces the viral RNA load of EBOV, MARV, and BDBV at low-nanomolar concentrations (**Figure 1C–G**). The calculated half-maximal and 90% inhibitory concentrations (IC□□ and IC□ □) for these filoviruses consistently ranged between 1.0 and 3.5 nM, highlighting zotatifin’s potent broad-spectrum activity. Importantly, this potent antiviral activity was achieved without inducing significant cytotoxicity at comparable doses (CC_50_ >200 nM) (**Figure S3B**), corresponding to a favorable predicted selectivity index (SI) >154. Selectivity Index 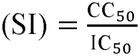

The clinical trial (<u>NCT04632381</u>) of zotatifin in non-hospitalized adults with mild or moderate COVID-19 included a pharmacokinetic (PK) component ^18,27^. Patients were randomized to a single subcutaneous dose of 0.01, 0.02, or 0.035 mg/kg or placebo. No patients developed Grade 3 or 4 adverse events. The relationship between drug exposure measures such as the time for which concentration exceeds the IC90 during the dosing interval (T>IC90), Cmax/IC90, AUC/IC90, and minimum concentration/IC_90_ (C_min_/IC_90_) achieved in patients were examined against the duration of viremia (based on swabs) using inhibitory sigmoid maximal effect (E_max_) models, with results shown in **Supplementary Table S1**. The PK/PD driver was %T>IC_90_, with an exposure mediating 50% of E_max_ (EC50) at a %T>IC90 of 42%, and the EC_80_ (target exposure) of 52.66% ±8.8% (r^2^ = 0.56). Given that the r^2^ was relatively low, we used machine learning approaches to identify predictors of duration of viremia. Gradient Boosting and Random Forest both identified T>IC_90_ among the top predictors of viremia duration. The first MARS basis function was %T>IC_90_ of 49% of the total dosing interval (MARS intercept = 15.492; generalized cross-validation = 19.290). These convergent estimates indicate that the target exposure is robust. This is the exposure that a dosing regimen must achieve in order to clear viremia. Because zotatifin’s mechanisms of action (eIF4A inhibition and induction of host immune responses) are expected to be shared between SARS-CoV-2 and filoviruses, we used a %T>IC90 of 50% as the exposure target for BDBV disease.

The zotatifin pharmacokinetic data from the PROPEL study were modeled using compartmental population PK analyses; the adult population PK parameters were used to model those in children.

The population PK parameter estimates for subcutaneous injection in 6 age groups are shown in **Table 1**.

**Table 1.** Zotatifin population pharmacokinetic parameter estimates across 6 age groups.

| **Age group** | **Weight** | ***CL*<sub>total</sub> L/h** | ***V<sub>c</sub>* L** | ***V<sub>p</sub>* L** | ***K<sub>a</sub>*h□¹** | ***CLD* L/h** |
|---|---|---|---|---|---|---|
| **Adult reference** | 80.46 kg | 17.30 ± 4.15 | 1300.0 ± 671.0 | 845.0 ± 23.2 | 19.60 ± 6.91 | 323.0 ± 12.8 |
| **Neonate, 1 month** | 4 kg | 0.69 ± 0.17 | 64.6 ± 33.4 | 42.0 ± 1.15 | 41.51 ± 14.64 | 34.0 ± 1.35 |
| **Infant, 1 year** | 10 kg | 2.107 ± 0.506 | 161.6 ± 83.4 | 105.0 ± 2.88 | 33.01 ± 11.61 | 67.6 ± 2.68 |
| **Child, 5 years** | 18 kg | 4.802 ± 1.152 | 290.8 ± 150.1 | 189.0 ± 5.19 | 28.46 ± 10.01 | 105.0 ± 4.16 |
| **Child, 12 years** | 40 kg | 9.450 ± 2.268 | 646.3 ± 333.6 | 420.1 ± 11.5 | 23.35 ± 8.24 | 191.1 ± 7.57 |
| **Adolescent, 16 years** | 60 kg | 12.968 ± 3.112 | 969.4 ± 500.0 | 630.1 ± 17.3 | 21.09 ± 7.43 | 258.4 ± 10.24 |

The large absorption rate constant (*K*_*a*_) means zotatifin is rapidly absorbed from subcutaneous tissue into the bloodstream, making it the best administration route. The between-patient variability in total clearance (CLtotal) of 24% is not clinically significant, with linear kinetics in the dose range tested. The large central volume (*V*_*c*_) means that there was extensive distribution into the central vascular compartment and highly perfused organs, the compartment in which BDBV viremia occurs^5,7^. Similarly, the large peripheral volume (*V*_*p*_) means that there was extensive distribution into infected peripheral organs. These results indicate extensive penetration of zotatifin into BDBV-infected tissues.

Monte Carlo experiments were used to identify the best dose for treatment of patients with symptomatic BDBV disease, in 540,000 virtual subjects across all age groups, following previously published methods ^28–30^. The dosing interval chosen was 2 weeks (336 h), in which six 2-week cumulative doses of 0.01, 0.02, 0.03, 0.1, 0.2, and 0.3 mg/kg were administered either as a single dose, or once weekly or daily for 7 days (that is, 0.0014, 0.003, 0.004, 0.014, 0.029, 0.043 mg/kg/day). The concentration-time profiles achieved in 5,000 virtual subjects for each of the 18 conditions in each of the 6 age groups are shown in Figures S4–S9. **Figure 1H** shows the proportion of 5,000 virtual patients treated with different daily doses who achieved or exceeded the target exposure (%T>IC90 = 50%) at an IC90 of 2.7 nM (0.0013 mg/L). The AUCs achieved in each age group are compared with those in the COVID-19 study and in the zotatifin oncology studies (NCT04092673) in Figure S10. Predicted AUCs were similar to those in the COVID-19 study, in which no patient developed Grade 3 or 4 adverse events, suggesting these doses and concentrations for BDBV treatment will also be well tolerated. Figures S4–S10 and Figure 1H show that the target exposure was attained at clinically tolerable doses in every age group, although not at the lowest doses simulated.

### Summary

Targeting host eIF4A-dependent translation offers a potential strategy for inhibiting genetically divergent filoviruses. Here, we identify purine-rich sequence features within ebolavirus 5′ UTRs and demonstrate potent in vitro antiviral activity of zotatifin against multiple filoviruses, including BDBV. Using an exposure target derived from patient-level pharmacokinetic– pharmacodynamic data (%T>IC90 ≥ 50%), Monte Carlo simulation in 540,000 virtual subjects across six pediatric and adult age groups identifies regimens that attain this target at exposures already tolerated in clinical trial participants. Together, these findings support further preclinical and translational evaluation of zotatifin as a host-directed antiviral candidate for filovirus disease. Its existing clinical development history provides a foundation for this work, but efficacy and safety in filovirus infection remain to be established.

## MATERIALS AND METHODS

Multiple sequence alignment and pairwise sequence identity calculations were performed using **Clustal Omega (version 2.1)**^**31–33**^ **with default parameters**.

### Heatmap of polypurine tract searching

5′ UTR sequences of seven Ebola virus protein-coding genes (NP, VP35, VP40, GP, VP30, VP24, and L) from four Ebola virus species: Ebola Zaire (NC_002549), Ebola Bundibugyo (NC_014373), SUDV Boniface (FJ968794.1), and TAFV (NC_014372) were retrieved from the NCBI (National Center for Biotechnology Information) GenBank and RefSeq databases.

For each 5′ UTR sequence, a 10-nucleotide sliding window was moved along the 5′ UTR sequence in single-nucleotide increments from the 5′ to the 3′ end, and the percentage of purine nucleotides (A+G) within each window was calculated as:

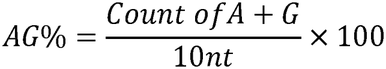

Results were visualized as a heatmap, with color intensity representing the local A+G percentage at each window position, ranging from red (low purine content, 0%) to green (high purine content, 100%). Windows exceeding 70% polypurine threshold are indicated by a red contour line. To enable comparison of polypurine tract localization across genes of different 5′ UTR lengths, each sequence was normalized to its total UTR length and displayed at relative positions from 0% (5′ end) to 100% (3′ end).

### Secondary structure prediction

Secondary structures of the 5′ UTR were predicted using RNAfold (v2.6.3) from the ViennaRNA package^34^. For each sequence, both the minimum free energy (MFE) and the thermodynamic ensemble were computed using the partition function algorithm, from which positional entropy was derived at each nucleotide position. MFE (expressed in kcal/mol) predicts the thermodynamic stability of a predicted RNA secondary structure, where more negative values signify higher thermodynamic stability. Additionally, the MFE frequency (%) denotes the statistical probability that the predicted secondary structure occurs within the full Boltzmann thermodynamic ensemble of all possible conformations.

### Population Pharmacokinetic Model and Parameter Selection

Individual zotatifin plasma concentrations were analyzed by compartmental modeling in ADAPT 5 (Biomedical Simulations Resource, University of Southern California) using the maximum-likelihood expectation-maximization (MLEM) algorithm, consistent with our published population PK and fractal-scaling analyses. Pediatric population PK parameter estimates were derived from the adult model using body-size scaling motivated by fractal geometry and an age-dependent maturation of CYP3A4-mediated oxidative dealkylation and subsequent glucuronidation, following published work on developmental PK and fractal variability ^35–39^.

### Monte Carlo experiments for dose finding

In silico dose-finding using Monte Carlo experiments (MCE) was performed for each of 6 doses and three dosing schedules (18 conditions), in 6 age groups, for 5,000 virtual subjects per condition (total of 540,000 virtual subjects)^40,41^. The design is a dose-fractionation study, shown in Table 2:

**Table 2.** Doses and dosing schedules evaluated in the Monte Carlo experiments.

| **Cumulative 2 week Dose (mg/kg)** | **Single dose schedule (mg/kg for each SQ injection)** | **Once a week schedule x 2 Doses (mg/kg for each SQ injection)** | **Once daily for 7 days (mg/kg for each SQ injection)** |
|---|---|---|---|
| **0.01** | 0.01 | 0.005 | 0.0014 |
| **0.02** | 0.02 | 0.01 | 0.0029 |
| **0.03** | 0.03 | 0.015 | 0.0043 |
| **0.1** | 0.1 | 0.05 | 0.014 |
| **0.2** | 0.2 | 0.1 | 0.029 |
| **0.3** | 0.3 | 0.15 | 0.043 |

The population PK estimates in **Table 1** were the domain of input entered into subroutine PRIOR. The probability of target attainment (PTA) is the proportion of the 5,000 virtual subjects in whom a given dose and dosing schedule (Table 2) achieves or exceeds the target exposure (%T>IC□ □ = 50%) at each IC□ □. For each virtual subject, concentration–time curves with 999 time slices were generated over a 2-week dosing interval (0–336 h). The dosing interval was chosen based on the duration of symptoms of 2-16 days in BDBV disease.

### Machine learning (ML) for efficacy in patients with COVID19

The predictor set included C_max_, C_min_, AUC_0–168_, AUC_0–336_, AUC to infinity, Absolute Time > IC_90_, exposure/IC_90_ ratios, %T> IC_90_, dose, age, sex, baseline weight, and baseline WHO score. Gradient boosting regression used shallow regression trees (maximum depth 2), Huber loss, shrinkage/learning rate 0.03, and deterministic random seed. Missing predictor values were median imputed within the analysis dataset. Predictor importance was based on tree impurity reduction. Candidate thresholds were extracted from high-gain tree splits. Five-fold shuffled cross-validation (reduced automatically where sample size required) was used to estimate MAE and R^2^.

MARS was implemented as a first-order adaptive hinge-basis regression: candidate basis functions took the form max(0, x−t) and max(0, t−x) at data-supported knots. Forward selection minimized residual sum of squares, followed by backward pruning using GCV. No interaction terms were permitted because of the small sample.

Random forest regression used bootstrap-aggregated regression trees, square-root feature sampling, minimum leaf size 2, and deterministic random seed. Predictor importance and highgain split thresholds were extracted. Cross-validated MAE and R^2^ were calculated using the same general resampling structure as for gradient boosting. Limited bootstrap refitting was used as a qualitative stability check of top predictors. Because 13 outcomes and numerous correlated predictors were explored without a prespecified multiplicity-controlled confirmatory hypothesis framework, no inferential p-values are presented and no threshold is designated as statistically validated.

### Detection and Quantification of EBOV, MARV, TAFV, and SUDV

All experiments involving EBOV, MARV, TAFV, SUDV were conducted at the BSL-4 facility of Philipps University of Marburg in accordance with national and international regulations.

### Cells and infections

HuH7 (human hepatoma cells, fully matching the short tandem repeat (STR) reference profile of cell line HuH7) and Vero C1008 (clone E6, VeroE6, ATCC CRL-1586, African green monkey kidney cells) cells were cultivated in Dulbecco’s modified Eagle medium (DMEM) supplemented with fetal bovine serum (10% for maintenance, 3% for experiments, 0% for infection), penicillin (50 units/ml) and streptomycin (50 µg/ml) (DMEM ++) at 37 °C and 5% CO_2_. MARV strain Musoke (GenBank accession number DQ217792), EBOV variant Mayinga (GenBank accession number NC_002549), SUDV strain Boniface (GenBank accession number FJ968794.1), TAFV (GenBank accession number NC_014372), and BDBV strain 200706291/Uganda (GenBank accession number MK_028856) were propagated in VeroE6 cells. Viral titers of each viral stock were calculated based on TCID50 or standard plaque assays^42^.

Cells, plated in a 96-well plate (1 × 10□ cells/well), were infected with the various viruses at a multiplicity of infection (MOI) of 0.1. The infectious supernatant was removed and replaced with fresh 3%++ medium containing zotatifin [in DMSO] as indicated. 10 μM remdesivir was added as a control. Cells were further incubated for 48 h at 37 °C and 5% CO2. CPE development was monitored by light microscopy using a LEICA DMi1.

### 50% Tissue Culture Infectious Dose (TCID_50_) assay

VeroE6 cells were cultured in 96-well plates to 50% confluence and infected with 10-fold serial dilutions of supernatants from infected cells. At 10 to 14 days post-infection (p.i.), when the cytopathic effect had stabilized, cells were analyzed by light microscopy. The TCID_50_/ml was calculated using the Spearman–Kärber method^42^.

### Sample collection for filovirus-specific RT-PCR

RNA extraction from supernatant of infected cells (EBOV, MARV, TAFV, SUDV) was performed using QiaAMP Viral RNA Kit (Qiagen) 48 h p.i. Briefly, 112 μl AVL buffer with 1.12 μl carrier RNA (both Qiagen) and 0.5 μl bromophenol blue per well were placed in a fresh 96-well plate. A volume of 28 μl of the cell supernatant was transferred to the inactivating AVL buffer, mixed and incubated for 10 minutes. 112 μl of pure ethanol (>99.8%, p.a.) were added.

To extract RNA from cell lysate, remaining supernatant was removed and 130 μl RLT-buffer (RNeasy Mini Kit, Qiagen) with 1.3 μl 2-Mercaptoethanol and 0.5 μl bromophenol blue were added. The plate was incubated for 10 minutes, followed by the addition of 130 μl 70% ethanol. Inactivated samples were transferred to a new 96-well plate and sealed with adhesive aluminum foil, after which they were discharged through the disinfecting immersion bath.

RNA was purified using a fully automatic nucleic acid extraction system (Tecan). Briefly, 200 μl sample was combined with 200 μl of DNA/RNA Shield in a deep-well plate and purified using the Quick-DNA/RNA viral MagBead kit (Zymo Research). The kit was used according to the manufacturer’s instructions with the following adaptations: 20 μl magnetic beads, 460 μl of wash solutions and 460 μl of ethanol were used and RNA was eluted from the magnetic beads by the addition of 50 μl DNase/RNase-Free Water.

### Filovirus-specific RT-PCR

Viral RNA isolated from supernatants was analyzed using RealStar® Filovirus Screen RT-PCR Kit 1.0 (for MARV, EBOV, SUDV, TAFV; targets L gene) (altona Diagnostics). The kits were used according to the manufacturer’s instructions with the following adaptations: 2.5 μl Master A and 7.5 μL Master B solution were mixed with 5 μl sample from RNA extraction. The internal control provided by the manufacturer was used for samples from the supernatant. A positive control RNA was incorporated into each plate. All RT-PCRs shown here were conducted for a total of 45 cycles.

### Cell viability assay

Cell viability was determined using the CellTiter-Glo® (Promega) according to the manufacturer’s instructions. Briefly, Huh7 cells were treated with zotatifin (0–200 nM) or a vehicle control (DMSO) for 48 hours. Relative luminescence units (RLU) were measured and normalized to the DMSO-treated control group to determine the percentage of viable cells.

Half-maximal cytotoxic concentrations (CC_50_) were determined from absolute IC50 values calculated using GraphPad Prism. Data were fitted to a four-parameter, variable-slope nonlinear regression model (log[inhibitor] vs. normalized response) with defined top, bottom, and baseline constraints where X represents the log(concentration).

### Immunofluorescence Assay to determine BDBV Infection Efficiency

All experiments with replication-competent BDBV were performed in the BSL4 laboratory of Texas Biomedical Research Institute (Texas Biomed). To test whether zotatifin affects BDBV replication, Huh7 cells were inoculated with the virus at a multiplicity of infection (MOI) of 0.1 for 2 h. The cells were washed with PBS, then overlaid with medium containing zotatifin at two-fold serial dilutions ranging from 6.25 nM to 0, equal concentrations of DMSO, or no treatment. After 48 h, the cell culture supernatants containing BDBV were titrated onto VeroE6 cells for 24 h, then fixed. All samples were stained with Hoechst 33342 dye to detect nuclei and sdNP antibody (generously provided by Andrew Hayhurst, Texas Biomed^43^) to detect infected cells. The samples were photographed using a Nikon automated system and analyzed by CellProfiler to quantify virus infection. The infection efficiency was calculated as the ratio of the number of BDBV-positive cells to the total number of Hoechst-stained nuclei^44^.

### Dose-Response Curve Fitting and Statistical Analyses

Dose-response curves were generated in GraphPad Prism (Version 11.0.2) using nonlinear regression analysis. Data were fitted to a four-parameter logistic model (log[inhibitor] vs. response - Variable slope) to determine the half-maximal inhibitory concentration (IC_50_) and Hillslope (h). To determine IC_90_ values and their confidence intervals, data were fitted to log(agonist) vs. response -- Find ECanything equation: 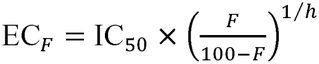 With F constrained to 10 (corresponding to 10% remaining activity) for a descending inhibition curve (h < 0).

## Acknowledgments

We are grateful to Andrew Hayhurst for providing sdNP antibody to detect BDBV infection. KR and SC would like to acknowledge the support from CRC1292.

## References

1. Rougeron, V., Feldmann, H., Grard, G., Becker, S. & Leroy, E. M. Ebola and Marburg haemorrhagic fever. J. Clin. Virol. 64, 111 (2015).

2. Mahanty, S. & Bray, M. Pathogenesis of filoviral haemorrhagic fevers. Lancet Infectious Diseases 4, 487–498 (2004).

3. Ebola disease caused by Bundibugyo virus, Democratic Republic of the Congo, Kenya and Nigeria. https://www.who.int/emergencies/disease-outbreak-news/item/2026-DON622.

4. Russell, N. et al. An urgent opportunity to ensure children’s access to oral post-exposure prophylaxis trials for Bundibugyo virus disease. Lancet Child Adolesc. Health 10, 780–781 (2026).

5. van der Ven, A. J. A. M., Netea, M. G., van der Meer, J. W. M. & de Mast, Q. Ebola Virus Disease has Features of Hemophagocytic Lymphohistiocytosis Syndrome. Front. Med. (Lausanne). 2, 4 (2015).

6. Rollin, P. E., Bausch, D. G. & Sanchez, A. Blood chemistry measurements and D-Dimer levels associated with fatal and nonfatal outcomes in humans infected with Sudan Ebola virus. J. Infect. Dis. 196 Suppl 2, (2007).

7. Liu, D. X. et al. Ebola Virus Disease Features Hemophagocytic Lymphohistiocytosis/Macrophage Activation Syndrome in the Rhesus Macaque Model. J. Infect. Dis. 228, 371 (2023).

8. Scotti, M. T. et al. Conserved Filovirus Proteins as Targets of Broad-Spectrum Antivirals. bioRxiv 2025.09.26.678902 (2025) doi:10.1101/2025.09.26.678902.

9. Jahrling, P. B. et al. Experimental infection of cynomolgus macaques with Ebola-Reston filoviruses from the 1989-1990 U.S. epizootic. Arch. Virol. Suppl. 11, 115–134 (1996).

10. Weik, M., Enterlein, S., Schlenz, K. & Mühlberger, E. The Ebola Virus Genomic Replication Promoter Is Bipartite and Follows the Rule of Six. J. Virol. 79, 10660 (2005).

11. Neumann, G., Watanabe, S. & Kawaoka, Y. Characterization of Ebolavirus Regulatory Genomic Regions. Virus Res. 144, 1 (2009).

12. Jain, S., Martynova, E., Rizvanov, A., Khaiboullina, S. & Baranwal, M. Structural and Functional Aspects of Ebola Virus Proteins. Pathogens 10, 1330 (2021).

13. Ernst, J. T. et al. Design of Development Candidate eFT226, a First in Class Inhibitor of Eukaryotic Initiation Factor 4A RNA Helicase. J. Med. Chem. 63, 5879–5955 (2020).

14. TaroncherLoldenburg, G. et al. Targeting the DEAD-Box RNA Helicase eIF4A with Rocaglates—A Pan-Antiviral Strategy for Minimizing the Impact of Future RNA Virus Pandemics. Microorganisms 9, 540 (2021).

15. Grentzmann, G., Ingram, J. A., Kelly, P. J., Gesteland, R. F. & Atkins, J. F. A dualluciferase reporter system for studying recoding signals. RNA 4, 479 (1998).

16. Biedenkopf, N. et al. The natural compound silvestrol is a potent inhibitor of Ebola virus replication. Antiviral Res. 137, 76–81 (2017).

17. Valdés-Torres, P. et al. Broad-spectrum antiviral activity of the synthetic rocaglate zotatifin against Mayaro virus and other viruses. Front. Cell. Infect. Microbiol. 16, 1752166 (2026).

18. aeFFECTOR Therapeutics Announces Positive Top-Line Results. https://www.globenewswire.com/news-release/2023/02/21/2612164/0/en/eFFECTOR-Therapeutics-Announces-Positive-Top-Line-Results-from-Phase-1b-Clinical-Trial-of-Zotatifin-for-the-Treatment-of-COVID.html.

19. Cencic, R. et al. Antitumor activity and mechanism of action of the cyclopenta[b]benzofuran, silvestrol. PLoS One 4, (2009).

20. Müller, C. et al. Broad-spectrum antiviral activity of the eIF4A inhibitor silvestrol against corona- and picornaviruses. Antiviral Res. 150, 123–129 (2018).

21. Peron, G. et al. Silvestrol, a potent anticancer agent with unfavourable pharmacokinetics: Current knowledge on its pharmacological properties and future directions for the development of novel drugs. Biomedicine and Pharmacotherapy 177, (2024).

22. Obermann, W. et al. Rocaglates as Antivirals: Comparing the Effects on Viral Resistance, Anti-Coronaviral Activity, RNA-Clamping on eIF4A and Immune Cell Toxicity. Viruses 14, 519 (2022).

23. eFFECTOR Therapeutics Announces Positive Top-Line Results from Phase 1b Clinical Trial of Zotatifin for the Treatment of COVID | FirstWord Pharma. https://firstwordpharma.com/story/5708018.

24. Sharma, M. et al. Phase 1/2 dose expansion study evaluating first-in-class eIF4A inhibitor zotatifin in patients with ER+ metastatic breast cancer. Journal of Clinical Oncology 41, 1080–1080 (2023).

25. Deflubé, L. R. et al. Ebolavirus polymerase uses an unconventional genome replication mechanism. Proc. Natl. Acad. Sci. U. S. A. 116, 8535–8543 (2019).

26. Gruber, A. R., Lorenz, R., Bernhart, S. H., Neuböck, R. & Hofacker, I. L. The Vienna RNA websuite. Nucleic Acids Res. 36, (2008).

27. Gordon, D. E. et al. A SARS-CoV-2 Protein Interaction Map Reveals Targets for Drug-Repurposing. Nature 583, 459 (2020).

28. Pasipanodya, J. & Gumbo, T. An oracle: antituberculosis pharmacokinetics-pharmacodynamics, clinical correlation, and clinical trial simulations to predict the future. Antimicrob. Agents Chemother. 55, 24–34 (2011).

29. Deshpande, D. et al. Ceftriaxone Efficacy for Mycobacterium avium Complex Lung Disease in the Hollow Fiber and Translation to Sustained Sputum Culture Conversion in Patients. J. Infect. Dis. 230, e230–e240 (2024).

30. Gumbo, T. et al. Selection of a moxifloxacin dose that suppresses drug resistance in Mycobacterium tuberculosis, by use of an in vitro pharmacodynamic infection model and mathematical modeling. J. Infect. Dis. 190, 1642–1651 (2004).

31. Goujon, M. et al. A new bioinformatics analysis tools framework at EMBL–EBI. Nucleic Acids Res. 38, W695–W699 (2010).

32. Madeira, F. et al. The EMBL-EBI Job Dispatcher sequence analysis tools framework in 2024. Nucleic Acids Res. 52, W521–W525 (2024).

33. Sievers, F. et al. Fast, scalable generation of high-quality protein multiple sequence alignments using Clustal Omega. Mol. Syst. Biol. 7, MSB201175-(2011).

34. Lorenz, R. et al. ViennaRNA Package 2.0. Algorithms Mol. Biol. 6, (2011).

35. Hope, W. W. et al. Population pharmacokinetics of micafungin in pediatric patients and implications for antifungal dosing. Antimicrob. Agents Chemother. 51, 3714–3719 (2007).

36. Rogers, Z. et al. The Non-Linear Child: Ontogeny, Isoniazid Concentration, and NAT2 Genotype Modulate Enzyme Reaction Kinetics and Metabolism. EBioMedicine 11, 118–126 (2016).

37. Jeena, P. M., Bishai, W. R., Pasipanodya, J. G. & Gumbo, T. In silico children and the glass mouse model: clinical trial simulations to identify and individualize optimal isoniazid doses in children with tuberculosis. Antimicrob. Agents Chemother. 55, 539–545 (2011).

38. Hiruy, H. et al. Subtherapeutic concentrations of first-line anti-TB drugs in South African children treated according to current guidelines: the PHATISA study. J. Antimicrob. Chemother. 70, 1115–1123 (2015).

39. Hall, R. G., Swancutt, M. A. & Gumbo, T. Fractal geometry and the pharmacometrics of micafungin in overweight, obese, and extremely obese people. Antimicrob. Agents Chemother. 55, 5107–5112 (2011).

40. Medicines Agency, E. Committee for Medicinal Products for Human Use (CHMP) Guideline on the use of pharmacokinetics and pharmacodynamics in the development of antimicrobial medicinal products. www.ema.europa.eu/contact (2016).

41. Ambrose, P. G. et al. Pharmacokinetics-pharmacodynamics of antimicrobial therapy: It’s not just for mice anymore. Clinical Infectious Diseases 44, 79–86 (2007).

42. Virology Methods Manual.

43. Sherwood, L. J. & Hayhurst, A. Generating Uniformly Cross-Reactive Ebolavirus spp. Anti-nucleoprotein Nanobodies to Facilitate Forward Capable Detection Strategies. ACS Infect. Dis. 8, 343–359 (2022).

44. Han, Z. et al. Compound FC-10696 Inhibits Egress of Marburg Virus. Antimicrob. Agents Chemother. 65, (2021).

