## Supplementary Figures for "Targeting host eIF4A with zotatifin inhibits Bundibugyo virus and other filoviruses"

**Figure S1**

**C**

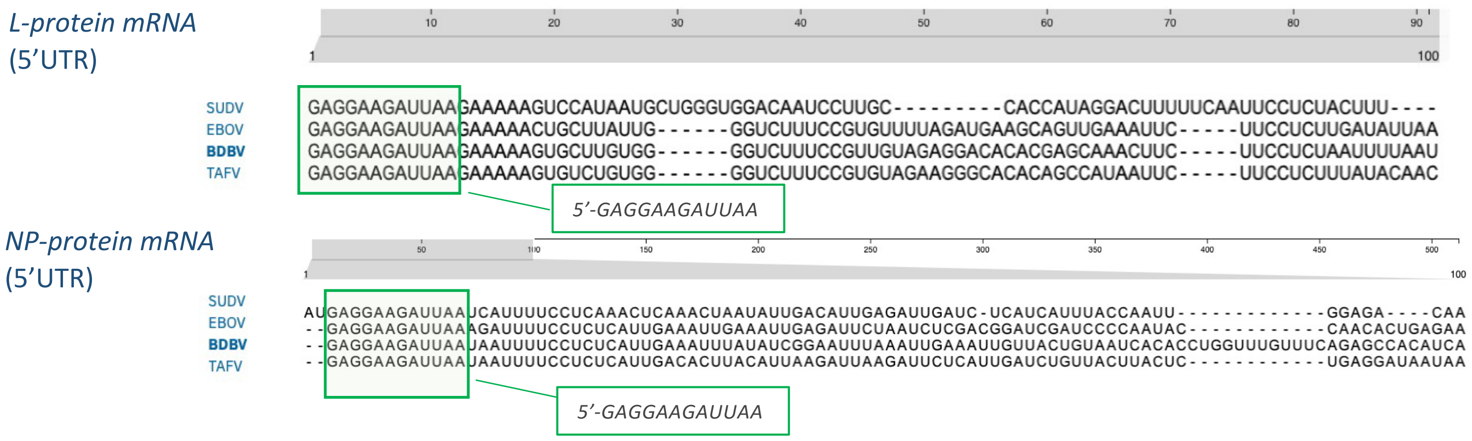


**Figure S1. Multiple-sequence alignment of 5′ UTRs of the primary structural (NP) and enzymatic (L) transcripts of four ebolavirus *species: Zaire (EBOV), Bundibugyo (BDBV), Sudan Boniface (SUDV) and Taï Forest (TAFV)*.** Nucleotide positions are indicated above each alignment, highlighting the pronounced length difference between the two transcripts. Polypurine-enriched, conserved nucleotide sequences at the most 5′-proximal positions are marked with green boxes; variable nucleotide positions across the viruses are shown, with alignment gaps represented by dashes.

**Figure S2**

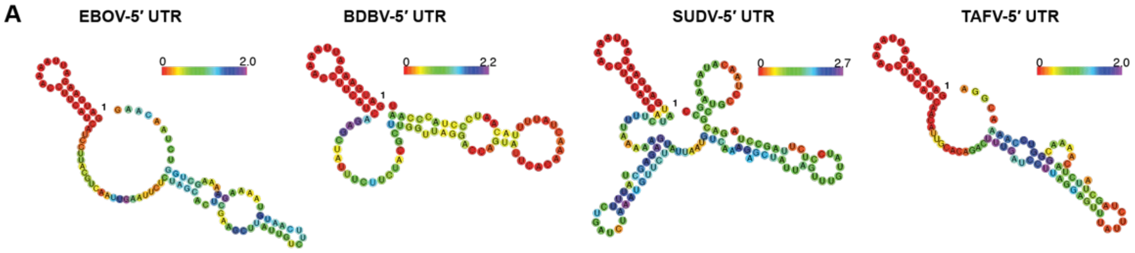


| | MFE (kcal/mol) | MFE Freq |
|---|---|---|
| EBOV | -15.90 | 11.9% |
| BDBV | -16.30 | 9.3% |
| SUDV | -23.50 | 2.0% |
| TAFV | -16.00 | 11.5% |

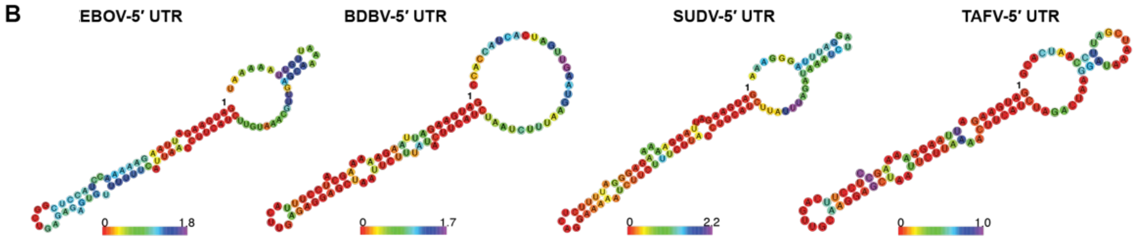


| | MFE (kcal/mol) | MFE Freq |
|---|---|---|
| EBOV | -19.90 | 4.4% |
| BDBV | -22.00 | 10.0% |
| SUDV | -22.70 | 3.4% |
| TAFV | -23.50 | 22.3% |

**Figure S2. The 5′-proximal regions of Ebola virus mRNA 5′ UTRs show conserved predicted secondary structures across four ebolavirus species.** Predicted secondary structures of the EBOV **(A)** VP35 and **(B)** VP40 gene 5′ UTRs generated by RNAfold. Shown is the minimum free energy (MFE) prediction with the corresponding MFE value and frequency in the summary tables. The color of each nucleotide indicates positional entropy.

**Figure S3**

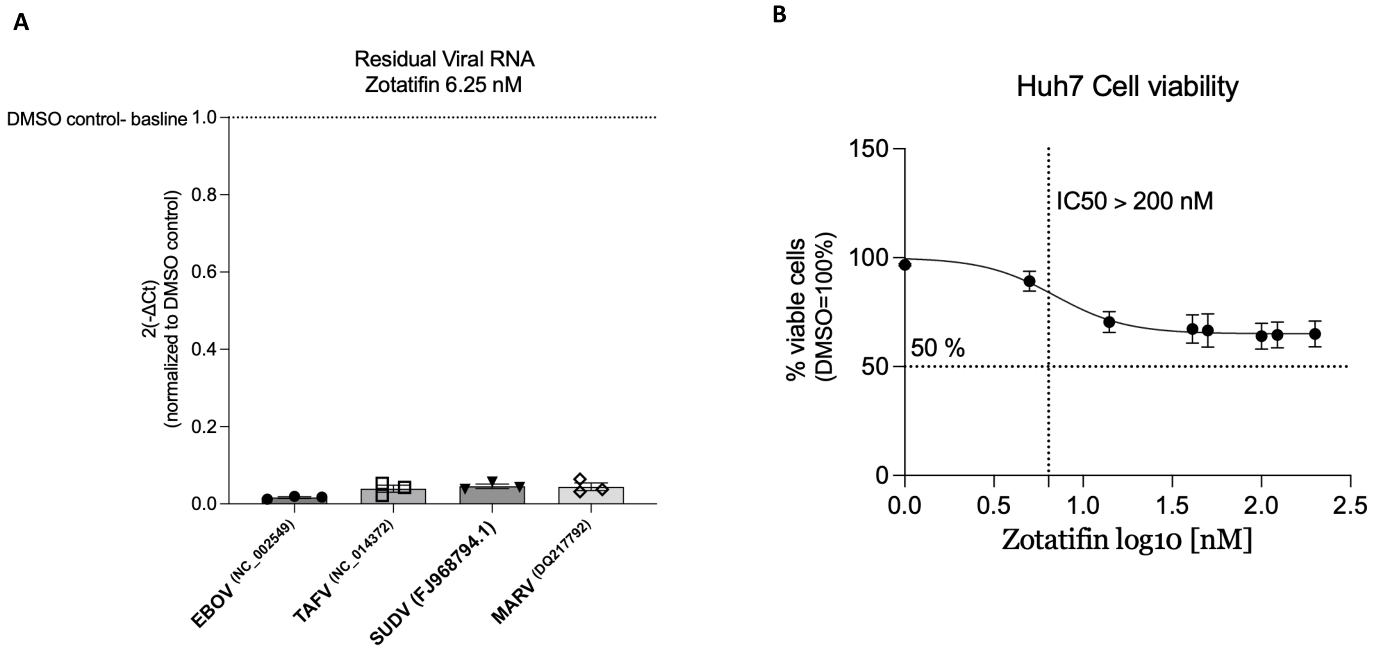


**Figure S3. (A) Viral RNA detected in supernatants of Huh7 cells infected with the indicated filovirus strain after treatment with 6.25 nM zotatifin.** Briefly, cells were infected at an MOI of 0.1 as indicated. After infection, zotatifin or vehicle control (DMSO) was added to the medium at concentrations ranging from 200 to 6.25 nM (two-fold dilutions). Supernatant from infected cells was collected 48 h post-infection, RNA was isolated and subjected to qRT-PCR detecting filovirus-specific RNA in the supernatant of infected cells. Shown are 2-( dCt value, normalized to DMSO control (1). Results are presented as mean ± SEM of at least three independent experiments (n = 3). **(B) Effect of zotatifin on Huh7 cell proliferation.** Briefly, Huh7 cells were treated with the indicated concentrations of zotatifin or vehicle control (DMSO) for 48 h. Cell viability was assessed using CellTiter-Glo® cell viability assay. Relative luminescence units (RLU) were normalized to DMSO-treated cells and are expressed as the percentage of viable cells. Results are presented as mean ± SEM of at least three independent experiments (n = 3). The $IC_{50}$values for prediction of the 50% cytotoxic concentration (CC50) were calculated using Prism software (log(inhibitor) vs. normalized response, variable slope, four parameters) and Absolute IC50, X is log(concentration) with defined top, bottom and baseline parameters.

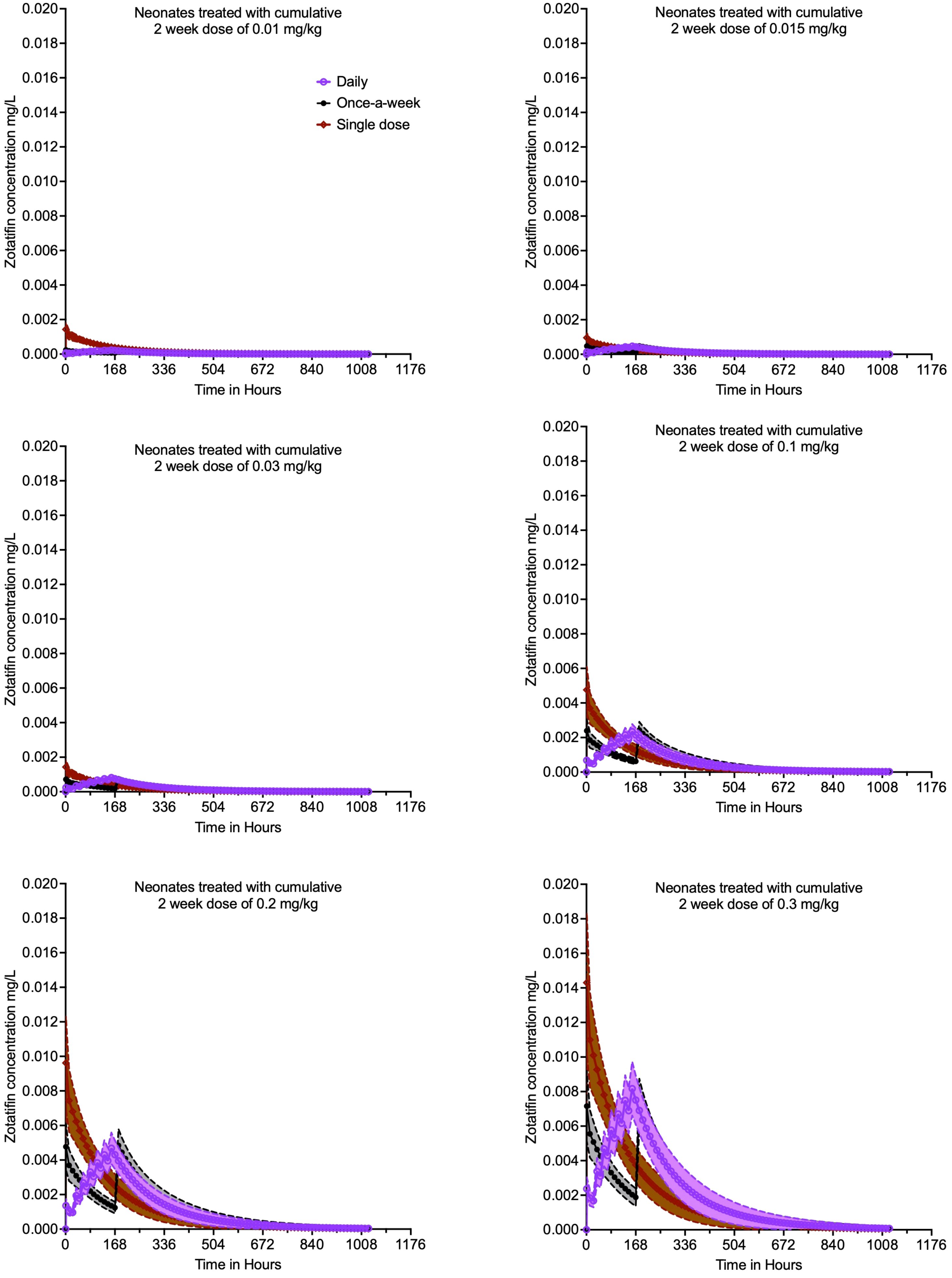


**Figure S4. Concentration-time profiles achieved in 30,000 virtual neonates.** In Monte Carlo experiments, 6 doses were administered to 5,000 patients for each of the three dosing schedules. Shown are median concentrations and 95% confidence intervals. The daily dosing shows accumulation of concentrations because of the long half-life.

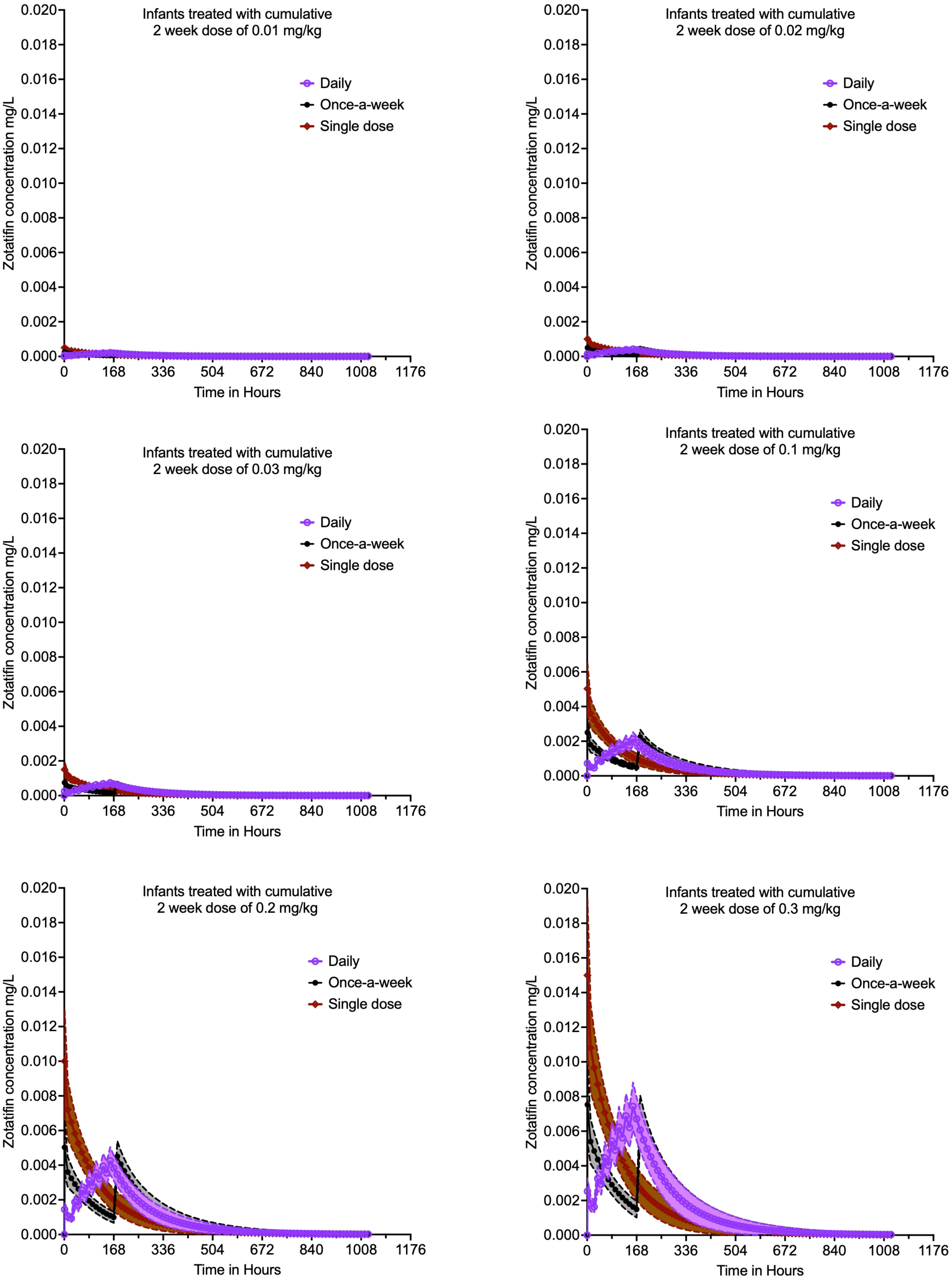


**Figure S5. Concentration-time profiles achieved in 30,000 virtual infants.** In Monte Carlo experiments, 6 doses were administered to 5,000 patients for each of the three dosing schedules. Shown are median concentrations and 95% confidence intervals. The daily dosing shows accumulation of concentrations because of the long half-life.

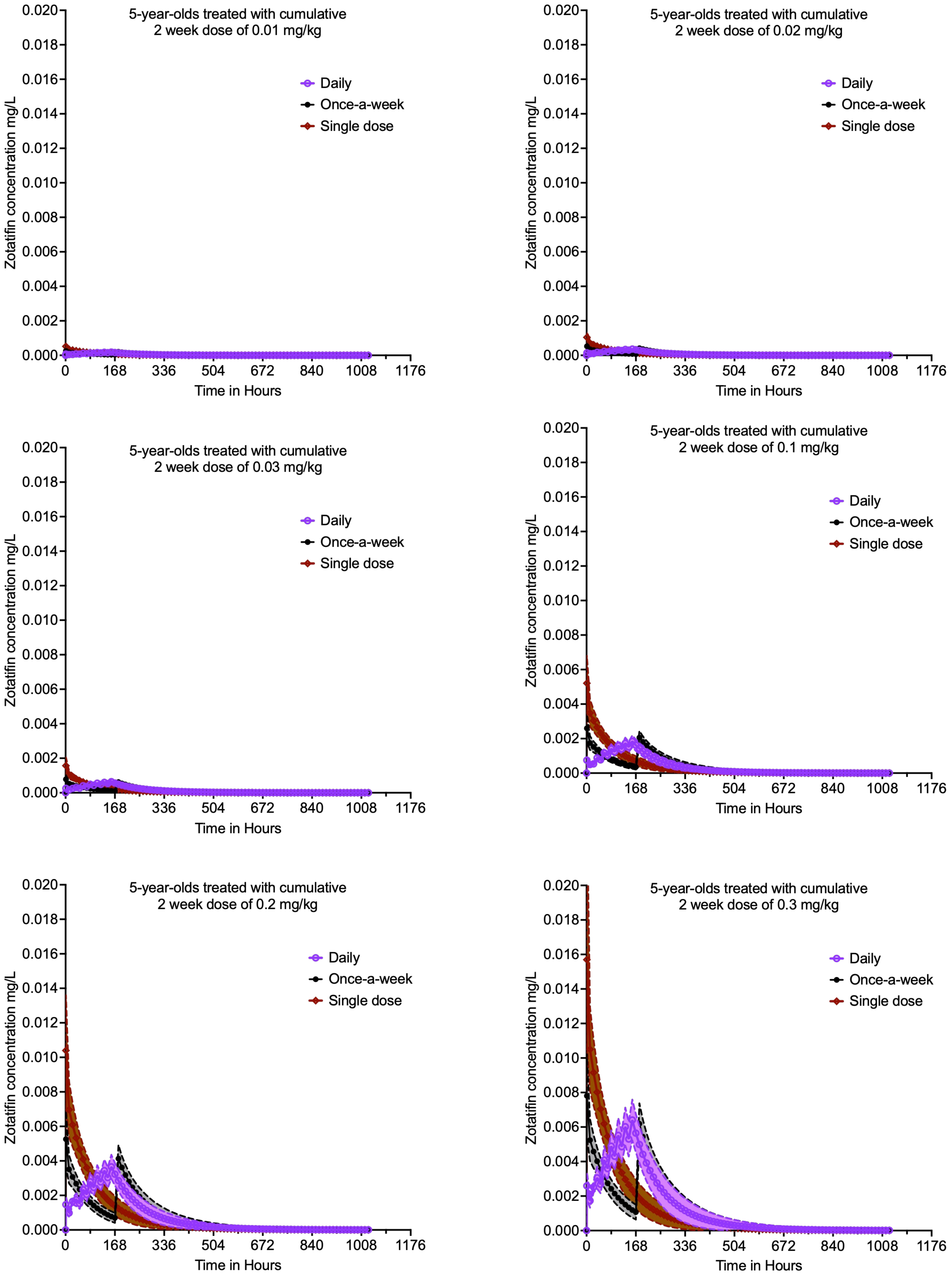


**Figure S6. Concentration-time profiles achieved in 30,000 5-year-old children.** In Monte Carlo experiments, 6 doses were administered to 5,000 patients for each of the three dosing schedules. Shown are median concentrations and 95% confidence intervals. The daily dosing shows accumulation of concentrations because of the long half-life.

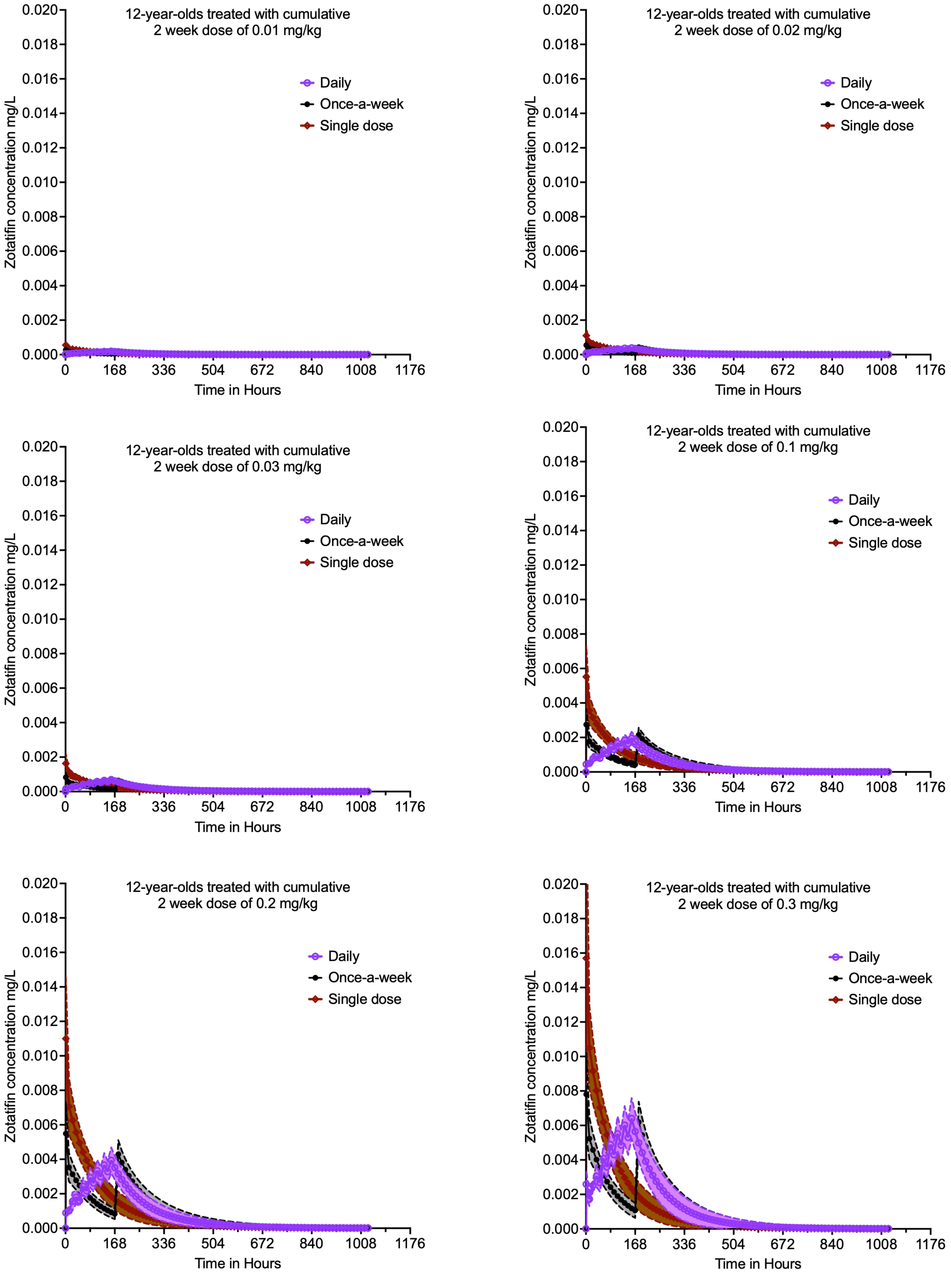


**Figure S7. Concentration-time profiles achieved in 30,000 12-year-old children.** In Monte Carlo experiments, 6 doses were administered to 5,000 patients for each of the three dosing schedules. Shown are median concentrations and 95% confidence intervals. The daily dosing shows accumulation of concentrations because of the long half-life.

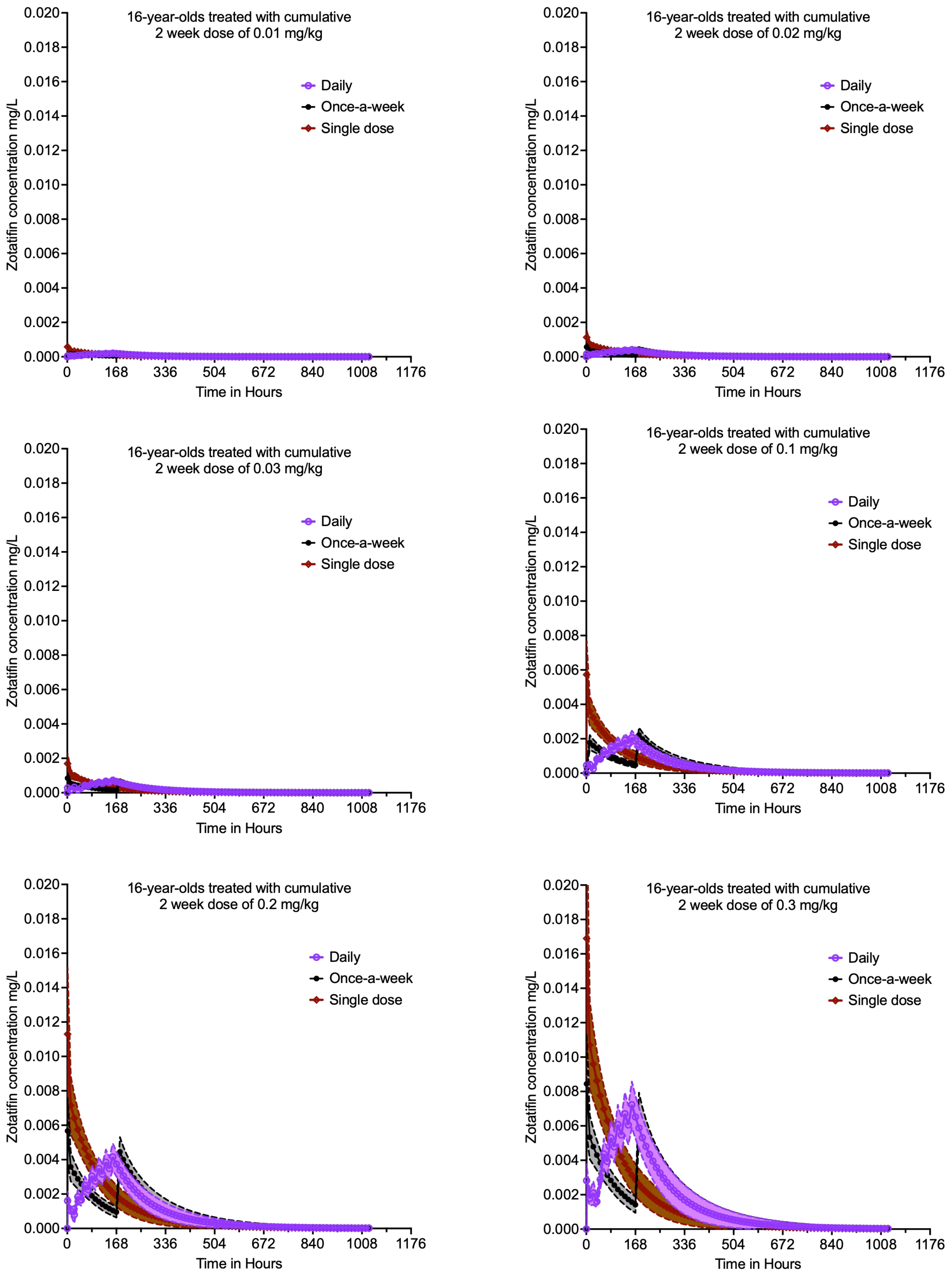


**Figure S8. Concentration-time profiles achieved in 30,000 16-year-old children.** In Monte Carlo experiments, 6 doses were administered to 5,000 patients for each of the three dosing schedules. Shown are median concentrations and 95% confidence intervals. The daily dosing shows accumulation of concentrations because of the long half-life.

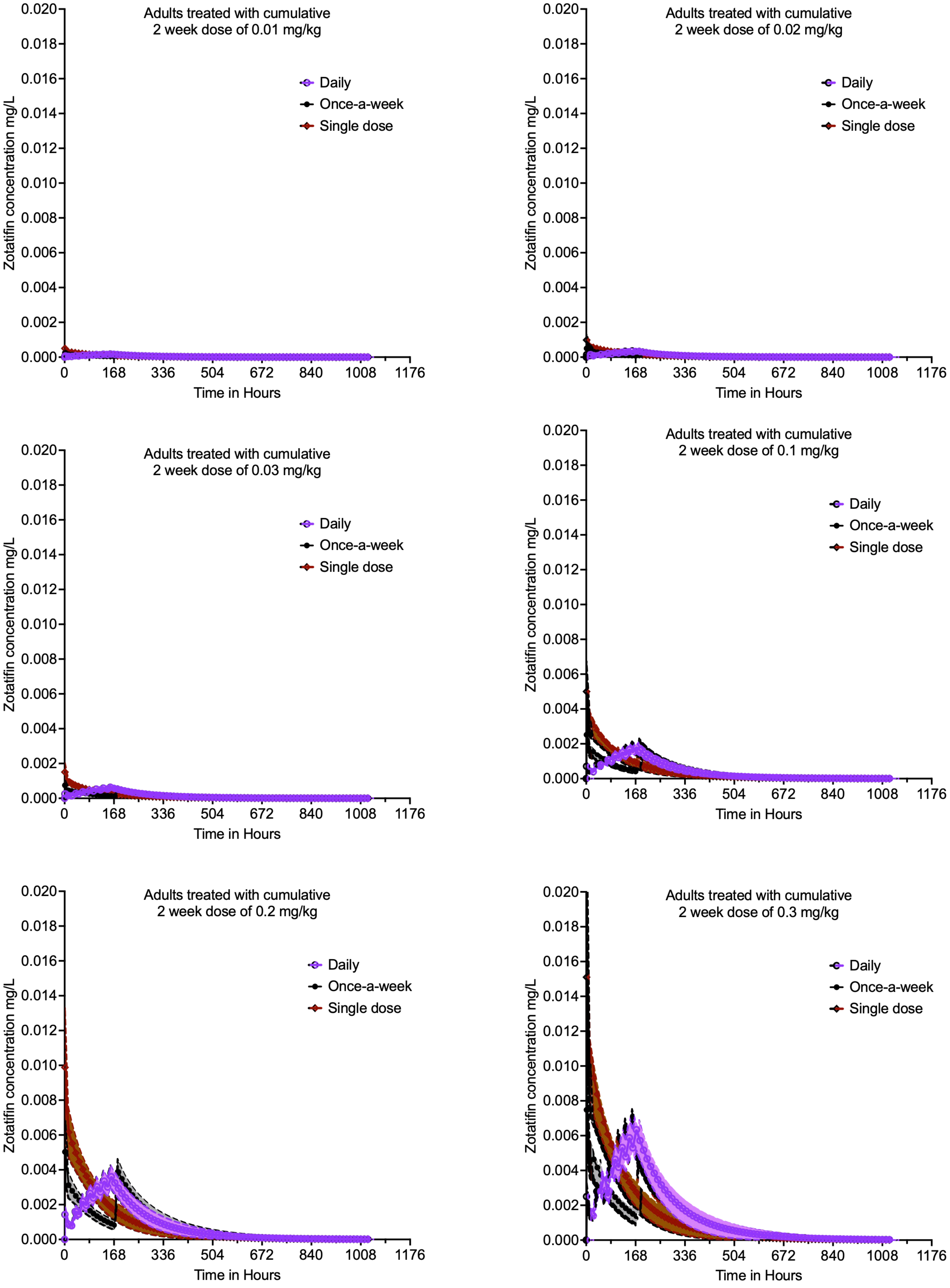


**Figure S9. Concentration-time profiles achieved in 30,000 adults.** In Monte Carlo experiments, 6 doses were administered to 5,000 patients for each of the three dosing schedules. Shown are median concentrations and 95% confidence intervals. The daily dosing shows accumulation of concentrations because of the long half-life.

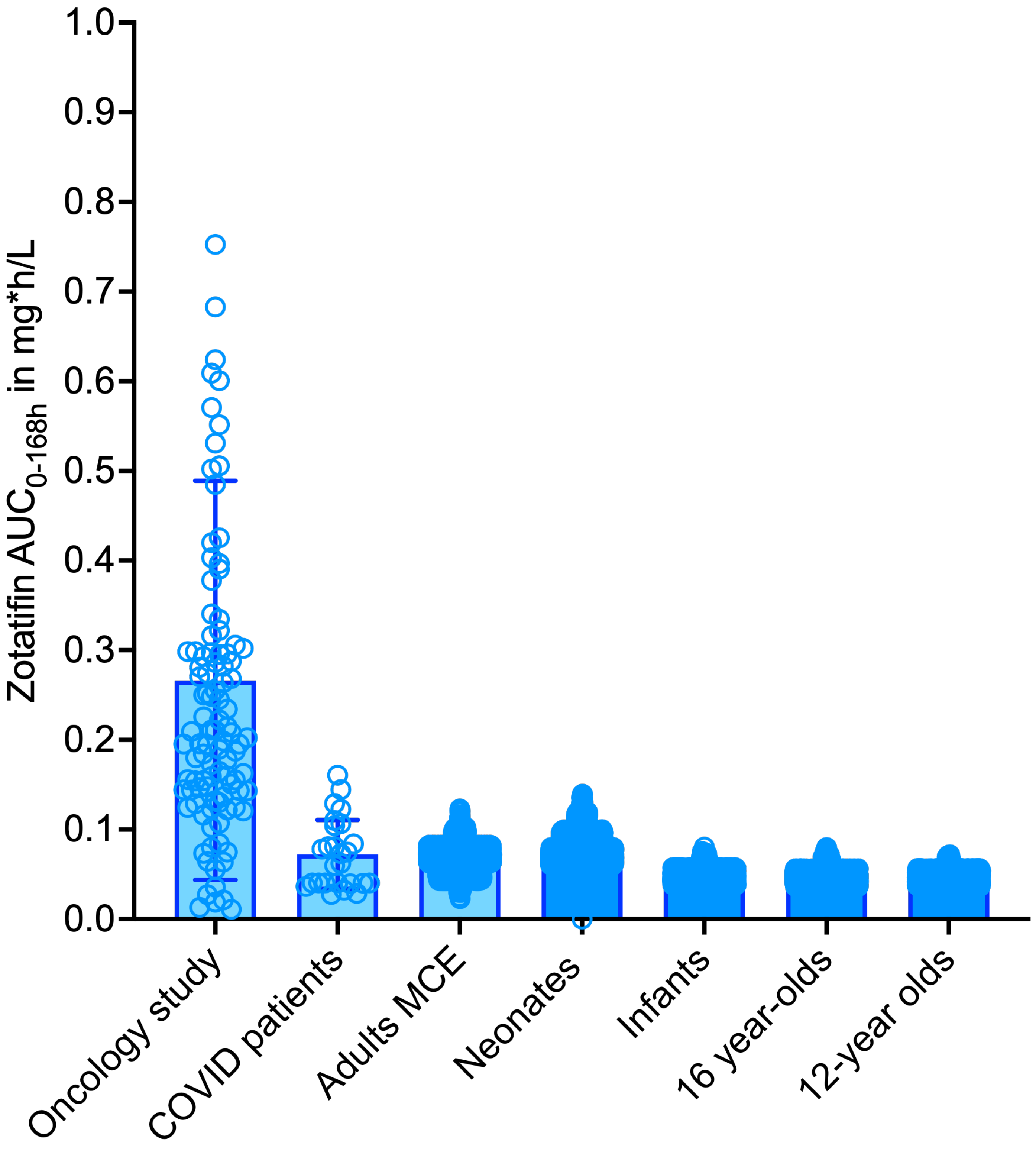


**Figure S10. AUC$_{0-168}$ at optimal dose in different age groups.** The AUCs were lower than in the oncology study and similar to those in the Covid19 study.
