## Supplementary table 1 for "Targeting host eIF4A with zotatifin inhibits Bundibugyo virus and other filoviruses"

**Supplementary Table S1. Inhibitory sigmoid E_max_ model-based comparisons of PK/PD drivers in COVID19 patients**

|  | **E_con_ (viremia days)** | **E_max_ (𝜟 days)** | **EC_50_** | **AICc** | **r^2^** |
| --- | --- | --- | --- | --- | --- |
| AUC_0-168_/IC_90_ | 11.34 | 23194.95 | NE* | 96.80 | 0.14 |
| AUC_0-336_/IC_90_ | NE* | NE* | NE* | 96.80 | 0.14 |
| C_max_/IC_90_ | NE* | NE* | NE* | 96.80 | 0.14 |
| C_min_/IC_90_ | NE* | NE* | NE* | 96.79 | 0.14 |
| %T>IC_90_ | 15 | 82.22 | 0.02 | 80.23 | 0.42 |
| T>IC_90_ | 15.01 | 16.77 | NE* | 80.52 | 0.41 |

*NE=not estimable because of poor convergence; %T>IC90= percentage of dosing interval (336h) that concentration persists above IC_90_; AICc=corrected Akaike Information Criteria score; AUC_0-168_= 0-168h area under the concentration-time curve; C_max_=peak concentration; C_min_=trough concentration; EC_50_ = pharmacokinetic-pharmacodynamic exposure mediating 50% of maximal effect; E_con_= viremia duration in placebo treated patient (days)s; E_max_= maximal effect in viremia duration (days); T>IC90= time in hours that concentration persists above IC_90_;
